# Cep57 and Cep57l1 cooperate to recruit the Cep63-Cep152 complex for centriole biogenesis

**DOI:** 10.1101/842682

**Authors:** Huijie Zhao, Sen Yang, Xiaomeng Duan, Qingxia Chen, Guoqing Li, Qiongping Huang, Xueliang Zhu, Xiumin Yan

## Abstract

Cep152 and Cep63 act as the cradle and recruit Plk4 to initiate the centriole biogenesis in a mother centriole dependent manner. However, how the Cep152-Cep63 complex is targeted to the proximal end of mother centrioles is unclear. In this study, we show that Cep57 and its paralog, Cep57l1, colocalize with Cep63 and Cep152 at the proximal end of mother centrioles in both cycling cells and differentiated multiciliated cells. Both Cep57 and Cep57l1 associate with the Cep152-Cep63 complex by directly binding to the centrosomal targeting region of Cep63. Co-depletion of Cep57 and Cep57l1 blocks the loading of Cep63-Cep152 to the centriole and subsequently prevents centriole duplication. We propose that Cep57 and Cep57l1 act together to ensure the recruitment of the Cep63-Cep152 complex to the mother centrioles for procentriole formation.

**Summary statement:** Cep57 and its paralog, Cep57l1, cooperatively act as a molecular scaffold to recruit the Cep63-Cep152 cradle to the proximal end of centrioles in the early stages of centriole duplication.

## Introduction

In mammalian cells, centrosomes function as spindle poles in mitosis and basal bodies for cilia formation. Typically, a centrosome is made of two cylindrical centrioles surrounded with pericentriolar material (PCM) (Nigg and Raff, 2009). In dividing animal cells, the duplication of centrioles must be tightly controlled to ensure the correct bipolar spindle formation, which is important for the integrity of genome. Aberration in centriole duplication is commonly present in numerous tumors and other diseases (Bettencourt-Dias et al., 2011; Gonczy, 2015; Nigg and Holland, 2018). Recent years, great progress has been made to understand the molecular mechanism of centriole biogenesis. To initiate the centriole duplication in mammals, Plk4 is firstly recruited by the Cep192-containing ring around the proximal mother centrioles at the early G1 phase and subsequently translocated to the larger Cep152-containing ring when Cep152 is recruited to the mother centriole(Kim et al., 2013; Sonnen et al., 2013). At the Cep152-ring, Plk4 recruits and phosphorylates Stil/Sas-5 to load Sas-6 for the cartwheel formation (Arquint et al., 2015; Arquint et al., 2012; Cizmecioglu et al., 2010; Dzhindzhev et al., 2010; Moyer et al., 2015; Ohta et al., 2014; Ohta et al., 2018). Several proteins, including Cep135, Cpap, Cp110 and Cep120, contribute to build the centriolar MT wall and mediate centriole elongation(Azimzadeh and Marshall, 2010; Brito et al., 2012; Carvalho-Santos et al., 2012; Comartin et al., 2013; Hung et al., 2004; Kohlmaier et al., 2009; Loncarek and Bettencourt-Dias, 2018; Schmidt et al., 2009). It is known that Cep152 is recruited by Cep63 to act as the cradle for the mother centriole dependent (MCD) centriole biogenesis (Brown et al., 2013; Kim et al., 2019; Sir et al., 2011; Zhao et al., 2013). However, it is still unclear how Cep63 or the Cep63-Cep152 complex is targeted at the proximal end of mother centrioles.

Cep57, a FGF-2 binding and trafficking protein, has been reported to be required for normal centriole duplication, proper chromosome segregation, PCM organization and centriole engagement (Bossard et al., 2003; Cuevas et al., 2013; Meunier et al., 2009; Snape et al., 2011; Wu et al., 2012; Zhou et al., 2016). Cep57, Cep63 and Cep152 form a stable complex at the proximal end of the centrioles and Cep57 is the proximity interactor of Cep63 (Firat-Karalar et al., 2014; Fortun et al., 2018; Lukinavicius et al., 2013; Sieben et al., 2018), suggesting that Cep57 may act as the upstream protein to target the Cep63-Cep152 complex for centriole duplication.

In this study, we find that Cep57l1 is the paralog of Cep57 and these two proteins recruit the Cep63-Cep152 complex for the centriole duplication.

## Results and Discussion

### Cep57 specifically co-localizes with Cep63 and Cep152 at the proximal end of mother centrioles

To investigate the spatial relationship of Cep57, Cep63 and Cep152, we costained Cep57 with either Cep63 or Cep152 in U2OS cells. Consistent with the previous study(Lukinavicius et al., 2013), 3D-structured illumination microscopy (3D-SIM) results showed that Cep57 formed a ring-like structure with Cep63 and Cep152 at the proximal end of both centrioles (Fig. 1A). Cep57 colocalized with Cep63 with a similar diameter (426.7 ± 49.7 nm for Cep57, 441.0 ± 39.6 nm for Cep63; 26 centrioles from 13 cells), while Cep152 stained a slightly bigger ring (511.0 ± 36.2 nm, 26 centrioles from 13 cells) (Fig. 1A,B). Consistently, GFP-Cep57 also enriched at the base of mother centrioles in U2OS cells. In addition to the centriolar localization, overexpression of GFP-Cep57 resulted in microtubule (MT) bundling and both Cep63 and Cep152 were recruited to the bundled MTs (Lin et al., 2013b; Momotani et al., 2008; Zhou et al., 2016) (Fig. 1C,D), indicating that Cep57, Cep63 and Cep152 form a complex and Cep57 could recruit Cep63 and Cep152 *in vivo*.

**Figure 1.**
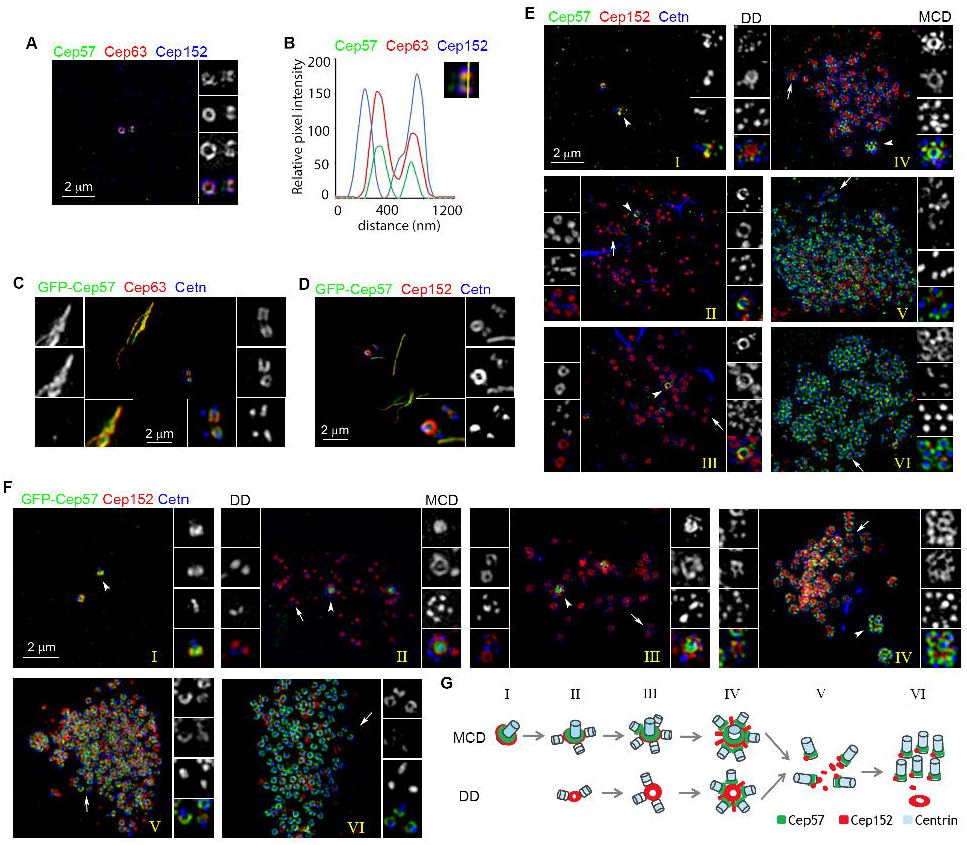
Cep57 co-localizes with Cep63 and Cep152 at the proximal end of centrioles. (A) Cep57 co-localized with Cep63 and Cep152. Cep57 (green) was co-stained with Cep63 (red) and Cep152 (blue) in human U2OS cells. (B) The line intensity plot showed the centriolar colocalization of Cep57, Cep63 and Cep152. (C, D) Cep63 and Cep152 were recruited to the bundled microtubule by GFP-Cep57 in U2OS cells. U2OS cells transfected with GFP-Cep57 for 48 hours were fixed and stained for Centrin (Cetn, blue) and Cep63 (C, red) or Cep152 (D, red). (E-G) Cep57 specified MCD cradles during centriole amplification in mTECs. mTECs (E) or mTECs expressing GFP-Cep57 (F) were fixed at day 3 after inducing multiciliation with the airliquid interfere (ALI d3) and stained for Cep152 (red), Cetn (blue) and Cep57 (green) or GFP. Cep152 serves as the marker of both the mother centriole and deuterosome, and Centrin as the marker of centrioles. Representative 3D-SIM images in the indicated stages of centriole amplification are presented. Magnified areas are used to show details for the MCD (right; arrowheads) and DD (left; arrows) centriole amplification. Schematic illustration models (G) are presented to aid understanding.

During multiciliation, mouse tracheal epithelial cells (mTECs) assemble hundreds of centrioles through both mother centriole-dependent (MCD) and deuterosome-dependent (DD) pathways respectively driven by Cep63 and its paralog, Deup1 (Lin et al., 2013a; Zhao et al., 2019; Zhao et al., 2013). At the early stages of centriole amplification, Cep63 specifically localizes at the proximal end of parental centrioles, but not at the deuterosomes, with the association of Cep152. Interestingly, Cep57 behaved similarly as Cep63 during mTECs differentiation (Fig. 1E,G). In stage I, Cep57 colocalized with Cep152 at the proximal end of two parental centrioles. In stages II and III, Cep57 exclusively associated with the parental centrioles, but not with the deuterosomes. In stage IV, Cep57 started to occupy the base of each procentriole formed around the parental centrioles and deuterosomes, where Cep152 started to protrude spoke-like structures. In stages V and VI, Cep57 still located at the proximal end of each nascent centriole as a full ring structure whereas Cep152 and Cep63 further converted into a dot (Fig. 1E) (Zhao et al., 2013). Consistent with the localization of endogenous Cep57, GFP-Cep57 exhibited a similar localization pattern in mTECs (Fig. 1F,G).

The spatiotemporal distribution of Cep57 in mTECs indicates that Cep57, similar as Cep63, only contributes to the MCD pathway and Cep57 together with Cep63 could recruit Cep152 for the MCD centriole assembly.

### Cep57, Cep63 and Cep152 interact hierarchically

To explore the relationship among Cep57, Cep63 and Cep152, we performed coimmunoprecipitation and GST pull-down experiments (Figs. 2, S1). Cep57 localized to the centriole via its C-terminal MT binding domain (Momotani et al., 2008) and interacted with Cep63 through its N-terminal coiled-coil domain (Figs. 2A and S1A,B). Cep63 bound to Cep57 via its N-terminal centriole-binding region and its C-terminus interacted with the centrosomal targeting region of Cep152 (152M) (Figs. 2B and S1C,D)(Hatch et al., 2010; Zhao et al., 2013). In the GST-pull down assay, GST-Cep57 directly interacted with His-Cep63, but could not pull down Cep152M in the absence of Cep63 (Fig. 2C), indicating that Cep57 forms the complex with Cep152 through Cep63 and these three proteins interact in a hierarchical manner (Fig. 2D). These results indicate that Cep57 localizes to the proximal mother centriole through its C-terminus and recruits the Cep63-Cep152 cradle by directly binding to the centriolar targeting region of Cep63.

**Figure 2.**
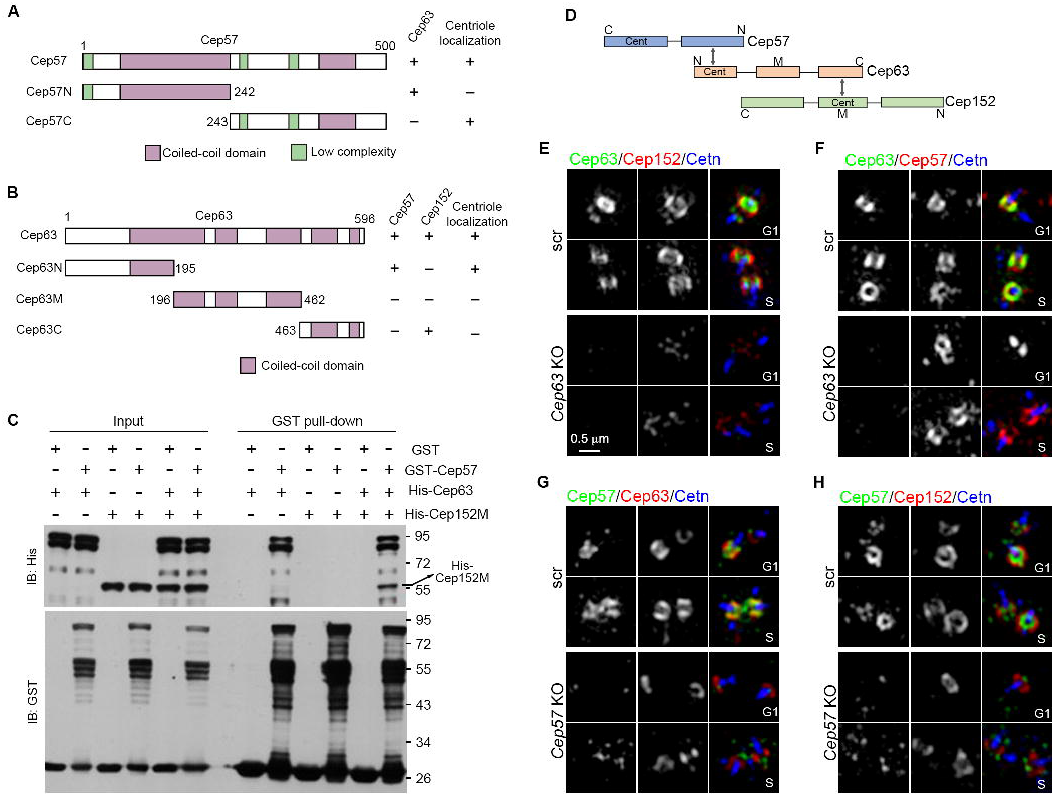
Cep57 is dispensable for anchoring the Cep63-Cep152 complex although directly interacting with Cep63. (A) Mapping the Cep63-interacting regions of Cep57. The numbers indicate amino acid positions. The N-terminus of Cep57 is required for the interaction with Cep63 and the C-terminus for its centrosomal localization. (B) Domain mapping of Cep63. The numbers indicate amino acid positions. The N-terminus of Cep63 is required for the interaction with Cep57 and its centrosomal localization, whereas the C-terminus interacts with Cep152 (Zhao et al., 2013). (C) GST-pull down assay. Bacterial lysates expressing the indicated proteins were mixed and subjected to GST pull-down assays. Cep152M contains the fragment of mouse Cep152 from positions 1075 to 1383 aa (Zhao et al., 2013). (D) Schematics showing the interaction order of Cep57, Cep63 and Cep152. Cent represents the centrosomal targeting region, and MT stands for the microtubule binding region. (E,F) The effects of knockout of *Cep63* on the centrosomal localization of Cep152 (E) or Cep57 (F) in U2OS cells. Note that knockout of *Cep63* abolished the centrosomal localization of Cep152, but not Cep57. (G,H) The effects of knockout of Cep57 on the centrosomal localization of Cep63 (G) or Cep152 (H) in U2OS cells. Note that both Cep63 and Cep152 were still localized to the centrioles upon the *Cep57* depletion.

### Depletion of Cep57 does not affect the centriolar localization of Cep63 and Cep152

To test whether Cep57 is required for the centrosomal targeting of Cep63 and Cep152, we generated *Cep57* and *Cep63* knockout (KO) U2OS cell lines using the CRISPR/Cas9 method (Fig. S2)(Cong et al., 2013). Immunofluorescence analysis of *Cep63* KO or *Cep57* KO cell lines revealed a complete loss of Cep63 or Cep57 ring-like signals at the proximal end of centrioles (Fig. 2E-H). As reported previously (Lukinavicius et al., 2013; Sir et al., 2011), Cep152 centrosomal signal reduced dramatically in *Cep63* KO cells compared to that in control cells (Fig. 2E). The centrosomal localization of Cep57 was not affected in *Cep63* KO cells (Fig. 2F). Surprisingly, both Cep63 and Cep152 were still able to localize at the centrosome in *Cep57* KO cells (Fig. 2G,H). These results are in accordance with the previous findings that Cep63 is essential for Cep152 centrosomal localization, whereas Cep57 seems to be dispensable for loading the Cep63-Cep152 cradle to the centrosome (Lukinavicius et al., 2013; Sir et al., 2011).

The dispensable role of Cep57 in the centriolar targeting of the Cep63-Cep152 complex is contradictory to the fact that Cep57 forms a stable complex with Cep63 and Cep152 by directly binding to the centriolar targeting region of Cep63. The paradox raises a possibility that other proteins may function redundantly with Cep57 to recruit Cep63 and Cep152.

### Cep57l1, the paralog of Cep57, forms a complex with Cep63 and Cep152 at the proximal end of mother centrioles

We searched for the homologous protein of Cep57 by protein BLAST and we indeed found out Cep57l1, the Cep57 paralog. Cep57l1 had similar function domains and shared approximately 43% identities with Cep57 (Figs. 3A and S3). We first tested whether Cep57l1 also interacts with Cep63 as Cep57. We found that the N-terminus of Cep57l1 was necessary and sufficient to bind Cep63 (Fig. 3B,C). Cep63 also interacted with Cep57l1 through its N-terminal centrosomal targeting fragment (Fig. 3E,F).

**Figure 3.**
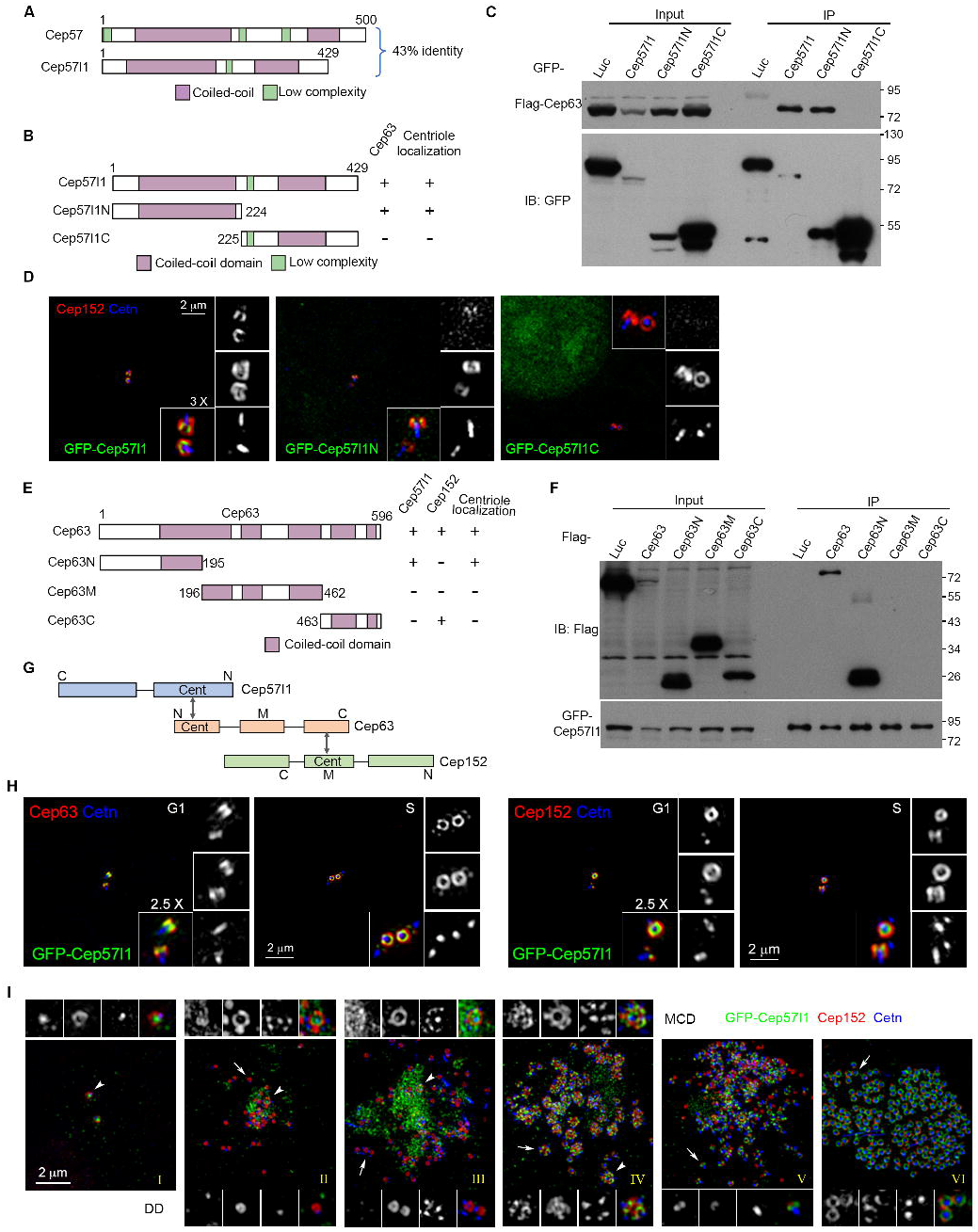
Cep57l1, the paralog of Cep57, forms a complex with Cep63 and Cep152 at the proximal end of centrioles. (A) Schematic diagram of mouse Cep57 and its paralog Cep57l1. (B) Schematic diagram of Cep57l1 and deletion mutants showing the ability to interact with Cep63 and to localize to the centriole. (C) Mapping the Cep63-interacting domain of Cep57l1. GFP tagged Cep57l1 or mutants were co-expressed with Flag-Cep63 in HEK 293T cells. Co-Immunoprecipitation was then performed with GFP-beads. (D) Identification of the centrosomal localization regions of Cep57l1. The N-terminus of Cep57l1 is required and sufficient for the centrosomal localization. U2OS cells were transfected with GFP-tagged Cep57l1 or mutants for 48h and the cells were stained for Cetn and Cep152. (E) Schematic diagram of Cep63 and deletion mutants showing the ability to interact with Cep57l1 and Cep152 and to localize to the centriole. (F) Mapping of the Cep57l1-interacting regions of Cep63. GFP tagged Cep57l1 was coexpressed with Flag tagged Cep63 or mutants in HEK 293T cells. Co-Immunoprecipitation was then performed with GFP-beads. (G) Schematics showing the interaction relationship of Cep57l1, Cep63 and Cep152. Cent represents the centriolar targeting region. (H) GFP-Cep57l1 co-localized with Cep63 and Cep152 at the proximal end of centrioles in U2OS cells. GFP-Cep57l1 was transiently expressed in human U2OS cells and co-stained with Cetn and Cep63 (left) or Cep152 (right). (I) The subcellular localization of GFP-Cep57l1 during centriole amplification in mTECs. Representative 3D-SIM images were acquired from mTECs at ALI d 3. Schematic illustration models are presented to aid understanding.

Due to the lack of the Cep57l1 antibody, we examined the subcellular localization of Cep57l1 by expressing GFP-Cep57l1 in U2OS cells. GFP-Cep57l1 also formed a ring-like structure with Cep63 and Cep152 at the proximal end of centrioles in the interphase with a comparable diameter to Cep57 (426.6± 34.2 nm, 40 centrioles from 24 cells) (Figs. 3H). Interestingly, although Cep57l1 showed high similarity with Cep57, Cep57l1 could not bundle microtubules (Fig. 3D,H). In addition, Cep57l1 was targeted to the centrioles through its N-terminal region (Fig. 3B,D), which was also different from Cep57 (Figs. 2A, S1A).

In multiciliated mTECs, GFP-Cep57l1 displayed similar subcellular distribution as Cep57 and Cep63 (Figs. 3I) (Lin et al., 2013a; Zhao et al., 2013). It also specifically associated with the parental centrioles but not with deuterosomes at the early stages of centriole amplification in mTECs (stages II and III). In stage IV, GFP-Cep57l1 started to accumulate at the base of nascent centrioles to form a goblet like structure (Fig. 3I). Like Cep57, GFP-Cep57l1 still localized at the proximal end of centrioles in a ring-like pattern in stages V and VI (Fig. 3I). Collectively, these findings imply that Cep57l1 is the paralog of Cep57 and like Cep57, Cep57l1 interacts with Cep63 at the proximal end of centrioles.

### Cep57 and its paralog Cep57l1 cooperate to recruit the Cep63-Cep152 complex for centriole duplication

To examine whether Cep57l1 is required for the localization of Cep63 and Cep152, we depleted Cep57l1 by RNAi in both wild type and *Cep57* KO U2OS cells (Fig. 4). Given the lack of Cep57l1 antibody, we selected Cep57l1 siRNA oligos by co-expression of GFP-Cep57l1 with various siRNA oligos and two oligos, 57L1i-1 and 57L1i-2, were chosen based on knockdown efficiency (Fig. S4).

**Figure 4.**
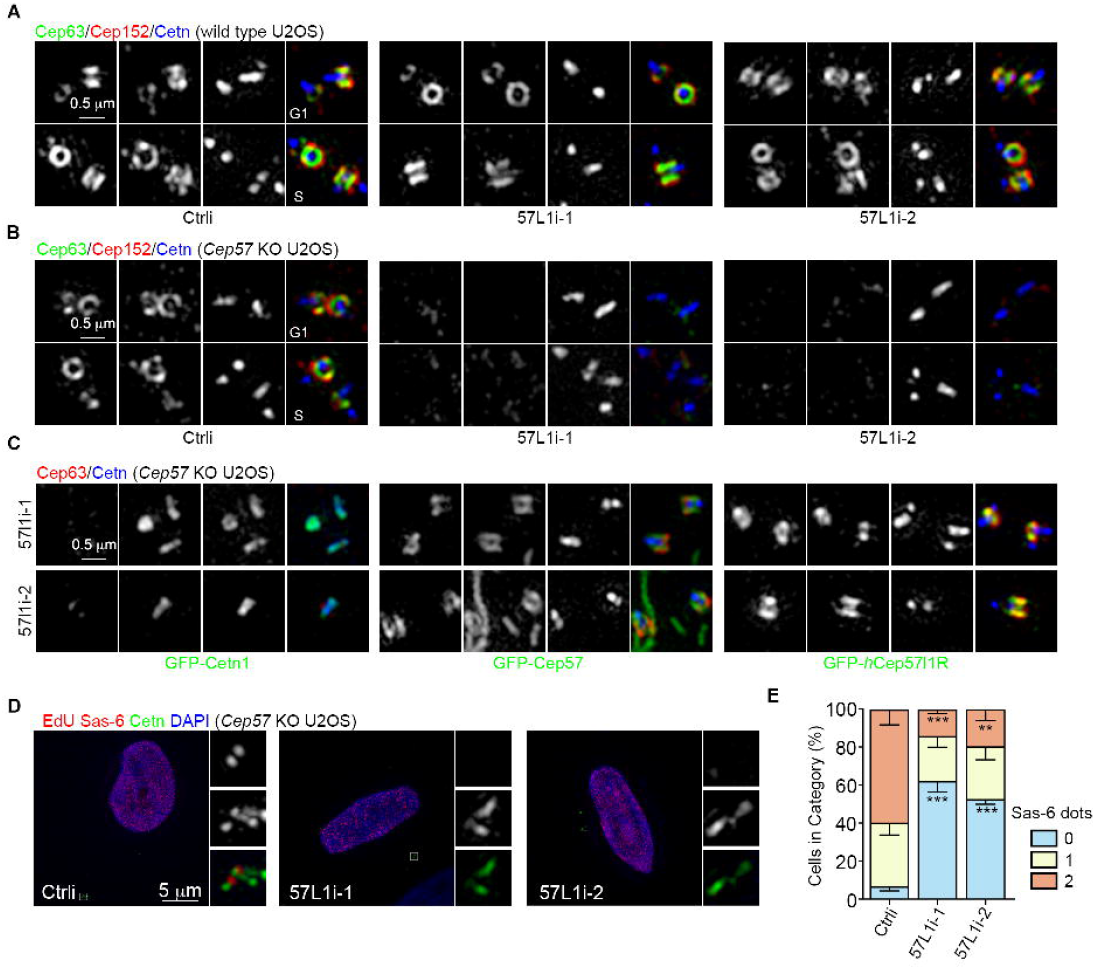
Cep57 and Cep57l1 recruit the Cep63-Cep152 cradle for centriole duplication. (A) Cep57l1 knockdown did not affect the centrosomal localization of Cep63 and Cep152. Human U2OS cells were transfected with a control siRNA or two Cep57l1 siRNA oligos respectively. (B) Double depletion of Cep57 and Cep57l1 abolished the centrosomal localization of Cep63 and Cep152. *Cep57* knockout cells were transfected with a control siRNA (Ctrli) or two Cep57l1 siRNAs (57L1i-1 and 57L1i-2) respectively. (C) Overexpression of both GFP-tagged Cep57 and RNAi-resistant *h*Cep57l1 fully rescued the centriolar loss of Cep63 upon the co-depletion of Cep57 and Cep57l1.For rescue experiments, *Cep57* KO U2OS cells were co-transfected with Cep57l1 siRNA oligos together with either GFP-Cetn1, GFP-Cep57 or RNAi resistant GFP-*h*Cep57l1 for 48 h and fixed for IF. (D) Co-depletion of Cep57 and Cep57l1 blocked procentriole formation. *Cep57* knockout cells were transfected with control or Cep57l1 siRNA oligos and stained for Sas-6 and Cetn. EdU was used to label S phase cells. (E) Percentage of cells with the indicated Sas-6 dots (n=100 cells). The results were from three independent experiments. Two tailed t-test was performed for p-values, error bars represent SD, and asterisks for p-values are ** p<0.01 and *** p<0.001.

As expected, like Cep57, depletion of Cep57l1 alone did not affect the centriolar localization of either Cep63 or Cep152 in control U2OS cells (Fig. 4A). In contrast, depletion of Cep57l1 in *Cep57* KO cells completely abolished the centriolar localization of Cep63 and Cep152 (Fig. 4B), suggesting that Cep57 and Cep57l1 act together to ensure the localization of the Cep63-Cep152 complex. Furthermore, the localization of Cep63 and Cep152 could be rescued by expressing either GFP-Cep57 or siRNA-resistant GFP-*h*Cep57l1 (GFP-hCep57l1R), but not GFP-Centrin1(GFP-Cetn1) (Fig. 4C).

To investigate whether procentriole assembly is impaired by co-depletion of both Cep57 and Cep57l1, we counted the number of assembled procentrioles marked by Sas-6 in S-phase cells. Consistent with the function of Cep63 and Cep152, the procentriole assembly was severely impaired upon the co-depletion of both Cep57 and Cep57l1 (Fig. 4D,E), demonstrating a critical function of Cep57 and Cep57l1 in the initiation of centriole duplication.

In this study, we find that Cep57 and its paralog Cep57l1 form a complex and colocalize with the Cep63-Cep152 cradle at the proximal end of mother centrioles. In multiciliated cells, similar as Cep63, Cep57 and Cep57l1 only associate with the parental centrioles but not with the deuterosomes at the early stages of centriole amplification, suggesting that Cep57, Cep57l1 together with Cep63 specifically function in the MCD pathway for recruiting Cep152. Co-depletion of Cep57 and Cep57l1 disrupts the localization of the Cep63-Cep152 complex at the mother centriole, sequentially blocks normal procentriole formation. Our results indicate that Cep57 and Cep57l1 cooperate to recruit the Cep63-Cep152 complex at the proximal end of mother centrioles for centriole biogenesis.

Recently, Cep295 is identified as the key protein for the conversion of daughter centrioles into functional mother centrioles, and the loading of Cep152 by Cep135 and Cep295 to daughter centrioles is critical for the conversion (Fu et al., 2016; Tsuchiya et al., 2016). Interestingly, Cep295 is only required for targeting of Cep152 to the young mother centriole, but not to the old mother centriole, implying that other scaffold proteins might play a role in anchoring Cep152 to the old mother centriole. Our findings here raise a possibility that Cep63 recruited by Cep57/Cep57l1 may function in detaining Cep152 at the mother centriole after its initial loading by Cep135-Cep295 during the centriole-to-centrosome conversion. In the future, it will be interesting to investigate the relationship among Cep57/Cep57l1, Cep63 and Cep295 in the centriolar localization of Cep152.

## Acknowledgements

The authors thank the Centre for Biological Imaging, Institute of Biophysics, CAS, for supports on 3D-SIM imaging, and institutional core facilities for cell biology and molecular biology for instrumental and technical supports. This work was supported by National Key R&D Program of China (2017YFA0503500) and Chinese Academy of Sciences (XDB19020000).

## Author contributions

X.Y. and X.Z. conceived and directed the project; H.Z. performed major experiments; S.Y. carried out quantification and generated knockout cell lines with the help of G.L., X.D. performed the GST pull-down experiment, Q.C. verified the localization of truncated proteins, and Q.H. assisted with the generation of the homemade antibodies; X.Y., X.Z., and H.Z. designed experiments, interpreted data, and wrote the paper.

## Additional information

The authors declare no competing interest.

## Supplementary Figure Legends

**Figure S1. Domain mapping of Cep57 and Cep63.**

(A) The C-terminus of Cep57 was required and sufficient for the centrosomal localization. U2OS cells expressing GFP tagged Cep57 or each fragment were co-stained with Cep152 (red) and Cetn (blue).

(B) Mapping the Cep63-interacting regions of Cep57. GFP-tagged Cep57 or mutants were co-expressed with Flag-Cep63 in HEK 293T cells for 48h and co-Immunoprecipitation was performed with GFP beads. The numbers indicate amino acid positions. The N-terminus of Cep57 was required for the interaction with Cep63 and the C-terminus for its centrosomal localization.

(C) The N-terminus of Cep63 was sufficient for the centrosomal localization. U2OS cells expressing GFP tagged Cep63 or each fragment were co-stained with Cep152 (red) and acetylated Tubulin (blue).

(D) Domain mapping of Cep63. GFP-tagged Cep57 was co-expressed with Flag-Cep63 or mutants in HEK 293T cells for 48h and co-Immunoprecipitation was performed with GFP beads. The numbers indicate amino acid positions. The N-terminus of Cep63 was required for the interaction with Cep57 and its centrosomal localization, whereas the C-terminus was required for interacting with Cep152.

**Figure S2. Sequencing confirmation of *Cep63* or *Cep57* knockout cell lines.**

**Figure S3. Sequence alignment of Cep57 and Cep57l1.**

The proteins sequence alignment was generated with the Clustal X 2.0 software using default parameters. The protein sequences were obtained from GenBank (*h*Cep57, NP_055494; *m*Cep57, NP_080941, *h*Cep57l1, NP_001077004 and *m*Cep57l1, NP_083408).

**Figure S4. Screening for *h*Cep57l1 siRNA oligos.**

HEK293T cells were co-transfected with GFP-*h*Cep57l1 and the indicated siRNA oligos for 48h and the cells were harvested for immunoblotting.

