## Supplementary material for "Cep57 and Cep57l1 cooperate to recruit the Cep63-Cep152 complex for centriole biogenesis": Materials and methods

**Cell culture, transfection and viral infection**

mTECs were isolated from 4-week C57BL/6J mice and cultured as described previously [1]. U2OS and HEK 293T cells were cultured in DMEM supplemented with 10% fetal bovine serum (FBS, ThermoFisher) at 37^o^C and 5% CO_2_. Cells were routinely tested for mycoplasma contamination by PCR. To observe the process of centriole amplification, mTECs were harvested at day 3 after the air-liquid interface (ALI) culture. For U2OS cells, plasmids were transfected using Lipofectamine 2000 according to the manufacturer’s instructions (ThermoFisher). For ectopic proteins expression and lentiviral production in HEK 293T cells, plasmids were transfected by the calcium phosphate method. Lentiviral infection of mTECs was performed as described previously [1]. Human CEP57L1 protein was depleted using two different siRNA duplex oligonucleotides that targeted the sequences 5’- CGTTGTACTCTTCTAGAGA -3’ (57L1i-1) and 5’- GCTTCAAACTGGACTTGAA -3’ (57L1i-2). A nontargeting siRNA duplex oligonucleotide was used as a control with the sequence 5’-TTCTCCGAACGTGTCACGTtt-3’ (Ctrli). U2OS cells were transfected with siRNA oligoes using Lipofectamine RNAiMAX reagent (ThermoFisher) for 48 hours. To improve the efficiency of knockdown, the second round of siRNA transfection was conducted for another 48 hours.

**Plasmids**

Full-length mouse cDNAs of Cep57 (accession no. NM_026665), Cep63 (accession no. NM_001301689), and Cep57l1 (accession no. NM_029132) were amplified from mTEC cDNA by PCR amplification. Full length human cDNAs of Cep57 (accession no. NM_014679) and Cep57l1 (accession no. NM_173830) were amplified from HEK 293T cDNA and the fragments were subcloned into pEGFP-C1. For antibody productions, full length of mouse Cep57 was then cloned into pGEX-4T-1. For ectopic expression, fragment of mouse Cep57, Cep63 and Cep57l1 were made by PCR amplification of the indicated regions and cloned into pEGFP-C1 or pcDNA3.1-NFLAG, respectively. For rescue experiments, RNAi-resistant GFP-Cep57l1 (GFP-*h*Cep57l1R) was made by site directed mutagenesis of pEGFP -Cep57l1. All constructs were verified by sequencing.

**GST pull-down assay**

GST and GST-Cep57 were purified with glutathione agarose beads (Sigma). His-Cep63 and His-Cep152M were purified with Ni-NTA agarose beads (QIAGEN). In vitro GST pull-down assays were performed by incubating recombinant proteins in lysis buffer (500mM NaCl, 50mM HEPES-KOH [PH 7.8], 5mM EDTA, 3mM DTT, 0.5mM PMSF and complete protease inhibitors (Chemicals)) for 4 hours at 4ºC. After three washes with lysis buffer, proteins were eluted with SDS-PAGE sample buffer and analyzed by SDS-PAGE.

**Generation of knockout cell lines**

The Cep57 and Cep63 knockout U2OS cell lines were generated by the CRISPR/Cas9 mediated gene editing system [2]. The sequences of sgRNAs were Cep57 (CACAAGCCCTAGCCATGCCG), Cep63 (AACATGCTGTAGAAGCTATA) and Scramble (GCACTACCAGAGCTAACTCA). The synthesized primers were annealed and inserted into pLV-Cas9- puromycin vector.

U2OS cells were transfected with the plasmids for 48 hours and were treated with 1.5μg/mL Puromycin (Sigma, P8833) for 10 days. The cells were then serially diluted and single colonies were examined by immunofluorescence microscopy to confirm the loss of the indicated proteins. Genomic DNAs isolated from the selected cells were PCR amplified with the following primers: Cep57-p (5’- GTGGTGTAGAGAATATAACTTCAAT-3’), Cep57-r (5’- CTCGATCATTGCACAAAGCTTTACT-3’), Cep63-p (5’- CT CATACATGAGGCCAGAATACAAA-3’) and Cep63-r (5’- GCCTAGTGCTTAATACCTAGT CATT-3’). The PCR products were subjected to sequencing.

**Immunoprecipitations**

Coimmunoprecipitation was performed as described previously[1]. In brief, HEK293T cells were transiently transfected, cultured for 48h, washed with PBS and lysed in lysis buffer (500mM NaCl, 50mM HEPES-KOH [PH 7.8], 5mM EDTA, 3mM DTT, 0.5mM PMSF and complete protease inhibitors (Chemicals)). Lysates were further cleared by centrifugation at 14000×g and subjected to coimmunoprecipitations using anti-Flag agarose beads (Sigma) or GFP-Trap agarose beads (Chromotek). Beads were washed in lysis buffer, eluted in sample buffer, and run on SDS-PAGE gels.

**Antibodies**

Primary antibodies for western blotting were directed against Flag (Sigma, F3165,1:5000), GFP (ThermoFisher, A6455, 1:3000), GST (Shanghai Wolwobiotech, Mab-GS01, 1:3000), and His (Sigma, H1029, 1:3000); Secondary antibodies were HRP-conjugated goat anti-Mouse IgG (H+L) (ThermoFisher, G21040, 1:5000) and goat anti-Rabbit IgG (H+L) (ThermoFisher, G21243,1:5000).

Primary antibodies purchased for immunostaining were directed against: GFP (ThermoFisher, A21311, 1:200), Centrin (Millipore, 04-1624, 1:200), Sas-6 (Santa Cruz, sc-81431, 1:50), Acetylated tubulin (Sigma, T6793, 1:1000), and Cep63 (Millipore, 2123674, 1:200). Chicken anti-Cep152 (1:300) and Rat anti-Cep57 (1:200) polyclonal antibodies were generated with purified GST fused proteins (Abclonal). The following secondary antibodies purchased from ThermoFisher were used for immunofluorescence: goat anti-Mouse IgG (H+L) Alexa Fluor 405 (A-31553,1:200) , goat anti-Rabbit IgG (H+L) Alexa Fluor 405 (A-31556,1:200), donkey anti-Rabbit IgG (H+L) Alexa Fluor 488 (A-21206,1:1000), goat anti-Chicken IgY Alexa Fluor 488 (A-11039,1:1000), goat anti-Chicken IgY Alexa Fluor 546 (A-11040,1:1000), goat anti-Rat IgG (H+L) Alexa Fluor 546 (A-11081,1:1000), and goat anti-Rabbit IgG (H+L) Alexa Fluor 546 (A-11035,1:1000).

**Fluorescence microscopy**

For immunofluorescence experiments, mTECs were pre-extracted with 0.5% Triton X-100 in PBS for 3 minutes before fixation with 4% fresh paraformaldehyde in PBS for 15 min at room temperature. For U2OS cells grown on coverslips, cells were pre-extracted with 0.5% Triton X-100 in PBS for 40 seconds and then fixed with 4% paraformaldehyde in PBS for 15 min at room temperature. After fixation, cells were permeabilized with 0.5% Triton X-100 in PBS for 15 min and blocked with blocking buffer (4% BSA in PBS) for 1 h. The incubation with primary and secondary antibodies was performed at room temperature for 2 h and 1 h, respectively. 3D-SIM images were collected on the DeltaVision OMX V3 imaging system (GE Healthcare). The setting of the OMX V3 system was as described previously[1].

**Statistics**

Results are presented as mean ± s.d. Two-tailed Student’s *t*-test (GraphPad Prism software) was used to calculate *P*-values between unpaired samples. Differences were considered significant when *P* <0.05.
