## Supplementary figures and images for "Cep57 and Cep57l1 cooperate to recruit the Cep63-Cep152 complex for centriole biogenesis"

### Supplemental Fig 1

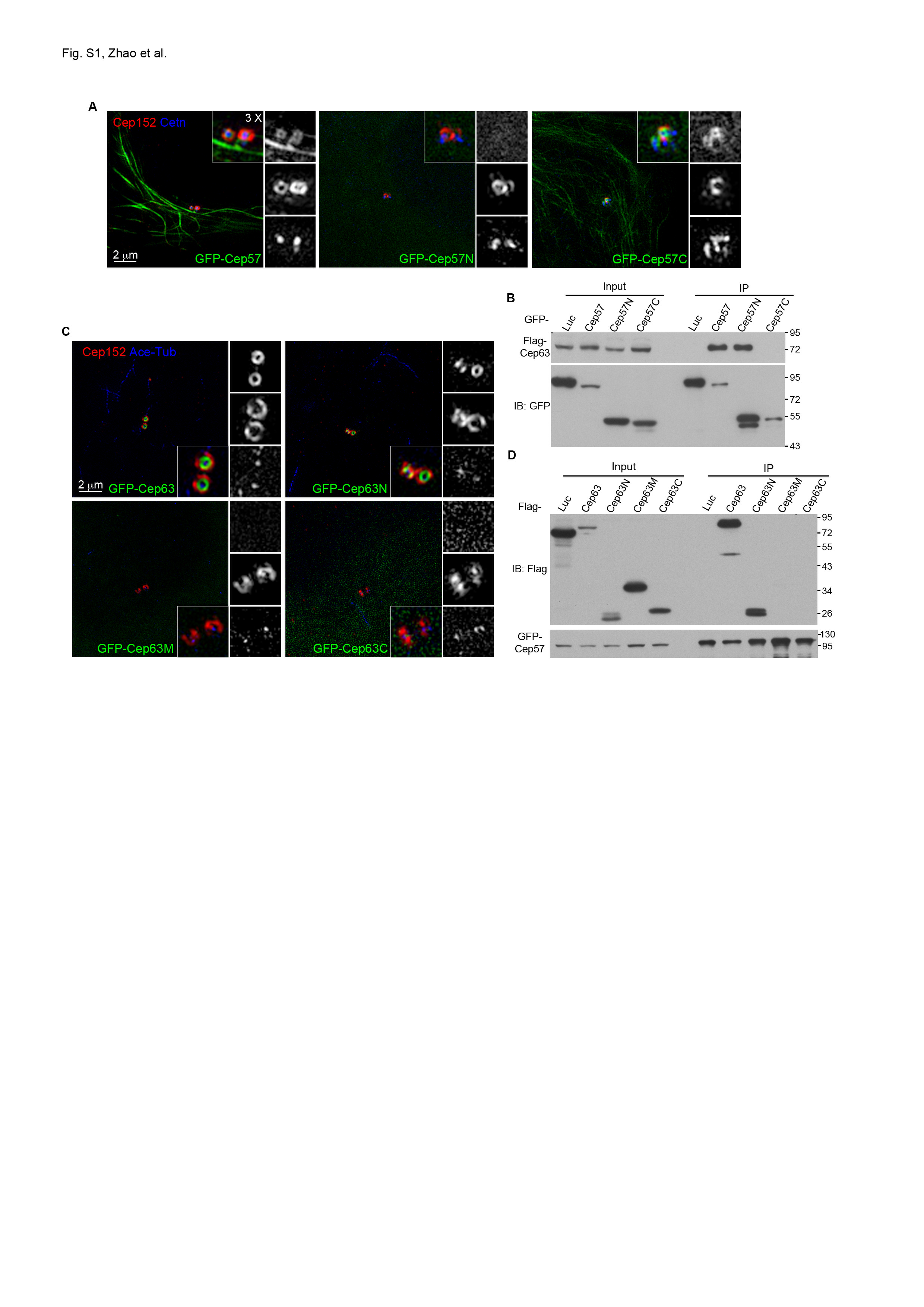

### Supplemental Fig 2

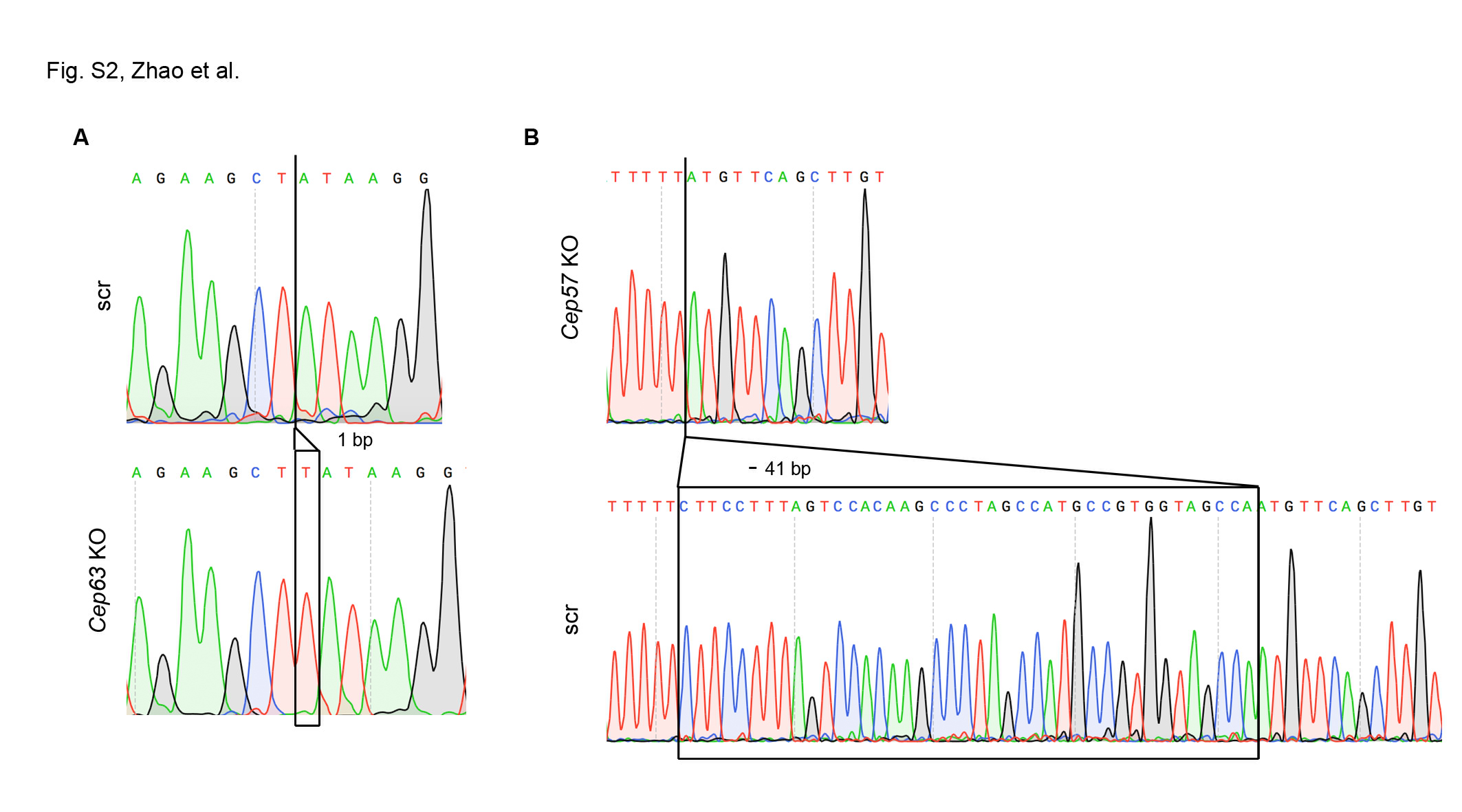

### Supplemental Fig 3

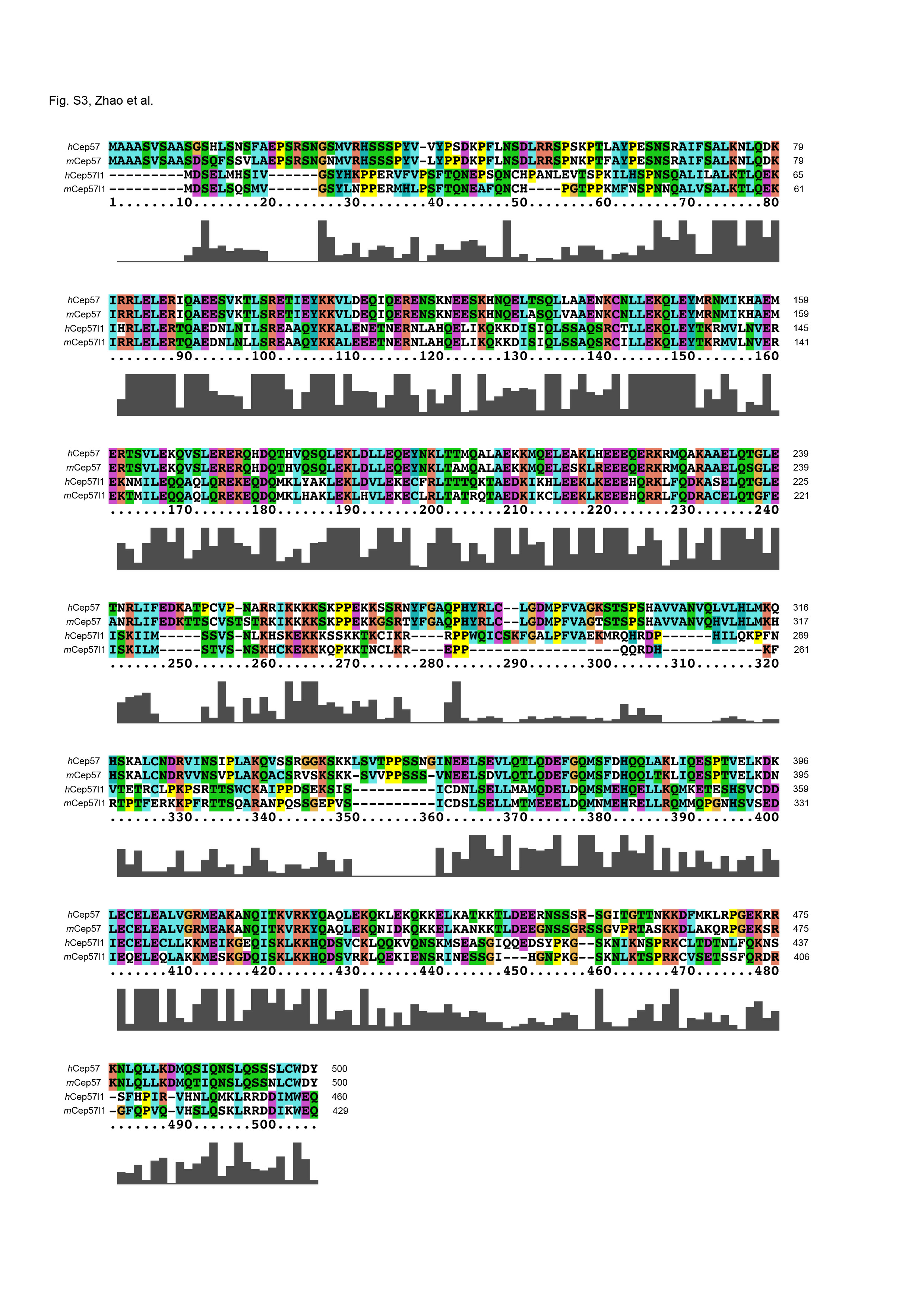

### Supplemental Fig 4

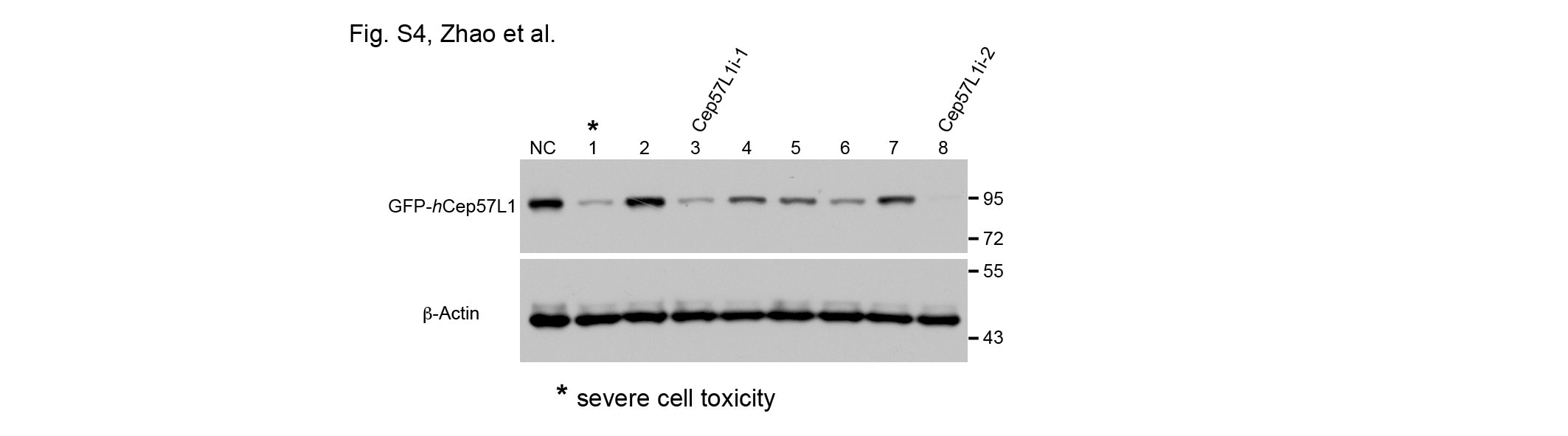
